# Outbreak of Glomerulonephritis by *Streptococcus zooepidemicus* SzPHV5 type, in Monte Santo de Minas, Minas Gerais, Brazil

**DOI:** 10.1101/330944

**Authors:** Rosângela S.L.A. Torres, Talita Z. Santos, Andre F. L. Bernardes, Patricia A. Soares, Ana C. C. Soares, Ricardo S. Dias

## Abstract

*Streptococcus zooepidemicus* is an emerging and opportunistic zoonotic pathogen, which playsan important role in the development of severe and life-threatening diseases and potentially capable of triggering large glomerulonephritis outbreaks. Between December 2012 and February 2013, 175 cases of glomerulonephritis were confirmed in the town of Monte Santo de Minas / MG / Brazil. During the outbreak, 19 isolates of *S. zooepidemicus* were recovered: one from ice cream, two from the oropharynx of food handlers and 16 from patients affected by acute post streptococcal glomerulonephritis (APSGN). All *S. zooepidemicus* involved in the outbreak amplified the same sequence of the hypervariable region of the SzP protein (SzPHV5) and presented indistinguishable banding patterns with high similarity (> 99%) by rep-PCR technique. Inspection programs on the milk supply chain should be strengthened and continuously encouraged so that consumers’ health is preserved.

---

***Streptococcus*** *equi* Subsp. *zooepidemicus* (Lancefield group C - GCS) is a common inhabitant of the respiratory and intestinal tract of horses, which, in special circumstances, behaves as a devastating and fatal pathogen for several animals (1). Humans (2) become infected when they get in contact with colonized or sick animals, or when they consume unpasteurized milk or its derived products (3,4). Infections caused by this microorganism have a wide spectrum of severity. They cause serious invasive diseases (bacteremia, septic arthritis, pneumonia and meningitis), posing a risk of death for the affected individuals (2,5), or still, benign diseases such as pharyngitis, but which can be followed by an episode of acute post streptococcal glomerulonephritis (APSGN) (4,6,7). APSGN is mediated by the deposition of immune complexes that lodge in the renal glomerulus after 7 to 21 days of primary streptococcal infection. The lesions resulting from this process lead to renal failure, albuminuria, hematuria, hypertension and edema (8,9). The disease is rare in industrialized nations; however, in the underprivileged world, new cases of APSGN range between 9.5 and 28.5 per 100,000 individuals per year (10).

**APSG**N is often triggered by the nephritogenic M-types of *Streptococcus pyogenes* (Lancefield group A-GAS) (7), or less frequently, by species of *S. zooepidemicus* e *S. disgalactiae* Subsp. *equisimilis* (Lancefield group C and G) (3,11,12).

***S. zooepidemicus*** present on their cell surface, a protein called SzP, which exhibits anti-phagocytic (M-like) properties similar to those found in EGA (13). The SzP protein varies the composition of the amino acids within the hypervariable region, as well as the amount of tetrapeptides (7 to 12-PEPK) of proline-glutamic acid-proline-lysine in the COOH-terminal region, determining 5 different types (HV1-HV5) (14). The hypervariable region of SzP can be used to genetically differentiate strains within the subspecies (5).

**Studies** have reported the use of repetitive element sequence-based PCR for the molecular typing of microorganisms. This method uses oligonucleotides complementary to repetitive highly conserved DNA sequences present at numerous copies in the bacterial genome. It enables genotypic characterization, clone differentiation, and clone dispersion in the community (15,16).

**Recently** we have described, in a control case study (17), a large outbreak of APSGN caused by *S. zooepidemicus,* in the small town of Monte Santo de Minas (population of 21.234 inhabitant), predominantly urban (77.4%), located in the south of Minas Gerais State. The cases were associated with the consumption of milk and its derived products. Forty-two patients (24.0%) were hospitalized and 4 individuals progressed to acute renal failure and required hemodialysis. No deaths were recorded.

**The** objective of this study is twofold. First, it aims to identify *S. zooepidemicus* strains collected from food handlers’ oropharynx secretion, and from milk, cheese and ice cream samples, during the glomerulonephritis outbreak in Monte Santo de Minas. Second, it attempts to genetically characterize the *S. zooepidemicus* strains isolated from the affected patients and from other sources during the outbreak, through the sequencing the hypervariable region of the SzP protein and the DNA fingerprint patterns with the DiversiLab^®^ software.

## Materials and Methods

### Study outline

**Between** December 25, 2012 and February 18, 2013, 417 suspected cases of APSGN were reported and 175 (42.0%) of which were confirmed by the Municipal Health Department of Monte Santo de Minas – MG.

**For** the microbiological identification of the etiological agent involved in the outbreak, 104 cow milk samples from several rural properties were collected; one sample from the mechanical milking machine; three from the storage tank; 25 samples of locally produced dairy-based foods products (cheese and ice cream); 11 samples of oropharyngeal secretions from food handlers who worked in 5 different ice cream parlors, 1 bakery and 1 dairy shop, located in different parts of the town; and 36 samples of oropharyngeal secretion from patients who developed pharyngitis and subsequent APSGN.

**The** samples were submitted for testing to the Laboratory of Microbiology of Foods - Ezequiel Dias Foundation, Central Laboratory of Minas Gerais (Lacen, MG). The samples of cow’s raw milk and isolates of group C beta-hemolytic streptococci identified in an ice cream sample, 2 handlers and 16 patients were tested at the Public Health Laboratory of the State of Paraná - Lacen/PR, Collaborating Center of the Ministry of Health-MS, Brazil, for the confirmation of species / subspecies and molecular characterization.

### Microbiologic Isolation and Identification

**Sample**s of raw milk were analyzed as previously published (4), with the following modifications: I-direct inoculation onto the surface of blood agar plates (BAPs); II-1 mL inoculation of the sample in 10 mL of Todd-Hewitt broth (BBL Microbiology Systems, Cockeysville, MD, EUA); III-1 mL inoculation of the sample in 10 mL of THB supplemented with nalidixic acid (15 mg / L) and colistin (10 mg/L). The cheese and ice cream samples were serially diluted in 0.1% buffered peptone water and then striated on the BAPs. Samples of oropharyngeal secretions from the patients and handlers, and from the milking machine teat cups were directly streaked out across on the BAPs.

**The** GCS isolates were presumptively identified based on their phenotypic characteristics on BAPs after incubation at 36°C±1 in aerobic atmosphere. The □hemolytic, catalase-negative, Gram-positive cocci arranged in chains, latex agglutination test group C – specific antisera (Streptococcal Grouping KitR, Oxoid, Basingstoke, England) were identified by the biochemical tests, deamination of arginine, hydrolysis of starch, production of acid in sorbitol, trehalose, lactose and ribose (18,19). Matrix-assisted laser desorption/ionization time-of-flight mass spectrometry (MALDI-TOF MS; Vitek MS, bioMérieux) was also employed to confirm the identify the strains involved in the outbreak (20).

### PCR and sequence analysis

***SzP*** gene was sequenced and the presumed protein sequence was determined for each isolate. PCR and DNA sequencing were performed as previously described (21), with PCR and sequencing primers cf1 (gataattaggagac atcatgtctagata), cf2 (ggctagcttcagtatcggcagccttgt), cr1 (aagctttaccactggggtat), and cr2 (gcaagagctgccgcggtgaa gaatggat) derived from the sequence with access number U04620 (21; bases 181 to 208, 274 to 300, 1362 to 1383, and 1276 to 1303, respectively) (22).

### Semi-automated rep-PCR DiversiLab®

**Genomic** DNA was extracted from a 10-μl loopful of *S. zooepidemicus* colonies using the UltraClean microbial DNA isolation kit (MO BIO Laboratories, Inc., Solana Beach, CA). PCR was performed using the DiversiLab *Streptococcus* kit (bioMérieux). PCR products were separated in a microfluidics DNA chip device (bioMérieux) in the Agilent 2100 BioAnalyser® (Agilent Technologies, Inc., Palo Alto, CA), according to the manufacturer’s recommendations. The relatedness was determined by cluster analysis and guidelines provided by the manufacturer. Isolates were categorized as indistinguishable, similar or different. In general, “different” was defined as ≤95% similarity and 2 or more band differences; “similar” was defined as ≥95% similarity and 1 band difference; and “indistinguishable” was defined as ≥95% similarity and no band differences (16).

### Statistics

**The** DNA fingerprint patterns were analyzed with the DiversiLab® software, 1.2.66 version (DiversiLab®, bioMerieux, France), using the Pearson correlation coefficient to determine distance matrices and unweighted pair group method with arithmetic mean (UPGMA) to create dendrograms, scatter plots, and electrophoregrams for data interpretation. Categorical variables were shown as frequencies and percentages.

### Results

***S. zooepidemicus*** was identified for presenting bacitracin resistance, negative CAMP-Test and Voges-Proskauer, positive arginine and esculin, and ability to ferment ribose, lactose, trehalose and sorbitol; slide agglutination group C; and confirmed by mass spectrometry (MALDI-ToF MS).

**The** samples obtained from the milk, storage tank and milking machine were negative for *S. zooepidemicus.* However, in 10 (9.6%) of the milk samples growth of *Streptococcus agalactiae* (Lancefield group B - GBS) was identified.

**Among** the 25 milk-derived food samples (cheese and ice cream) analyzed, only 1 (4.0%) ice cream sample, collected in one of the APSGN patient’s freezer, was positive for *S. zooepidemicus.*

**Among** 175 confirmmed APSGN patients, 36 oropharyngeal samples were analised, 16 (44.4%) presented positive culture test. Many culture negative patients had previously been treated with antimicrobials. Among the 11 oropharyngeal samples collected from food handlers, 2 (18.2%) were positive for *S. zooepidemicus,* one was an asymptomatic carrier and the other had pharyngitis (fever, swallowing pain, hyperemia, edema, and a pus-filled spot on one of the tonsils) and developed APSGN. One (9.1%) food handler was identified as carrying EGA *emm*-type 3. All patients involved in the outbreak were treated with 500 mg amoxicillin every eight hours for ten days. After the end of the antibiotic therapy, a new collection of oropharyngeal secretion was performed. All cultures were negative for *S. zooepidemicus*

**The** chromosomal templates from the 19 outbreak/2012-2013/Monte Santo-MG-Brazil isolates of *S. zooepidemicus* amplified the *szp* gene as expected and all PCR products shared sequence identity across their entire length (1,128-bp structural gene plus 19 bp of upstream sequence). These isolates were compared with strains of *szp* HV5 *-* 5058-APSGN /Nova Serrana-MG, Brazil/1998 (*22*) and revealed 100% identity with the 183 bases of the HV region.

**Highly** preserved regions of bacterial DNA of the 19 isolates of *S. zooepidemicus* were amplified to create a matrix of genetic proximity. Molecular typing (rep-PCR) produced a unique electrophoretic pattern (G1) of high genetic similarity (greater than ≥99%) and no different band was identified among the *S. zooepidemicus* from different sites, oropharynx of patients, food and ice cream handlers (**Figure 1**).

**FIG.1.**
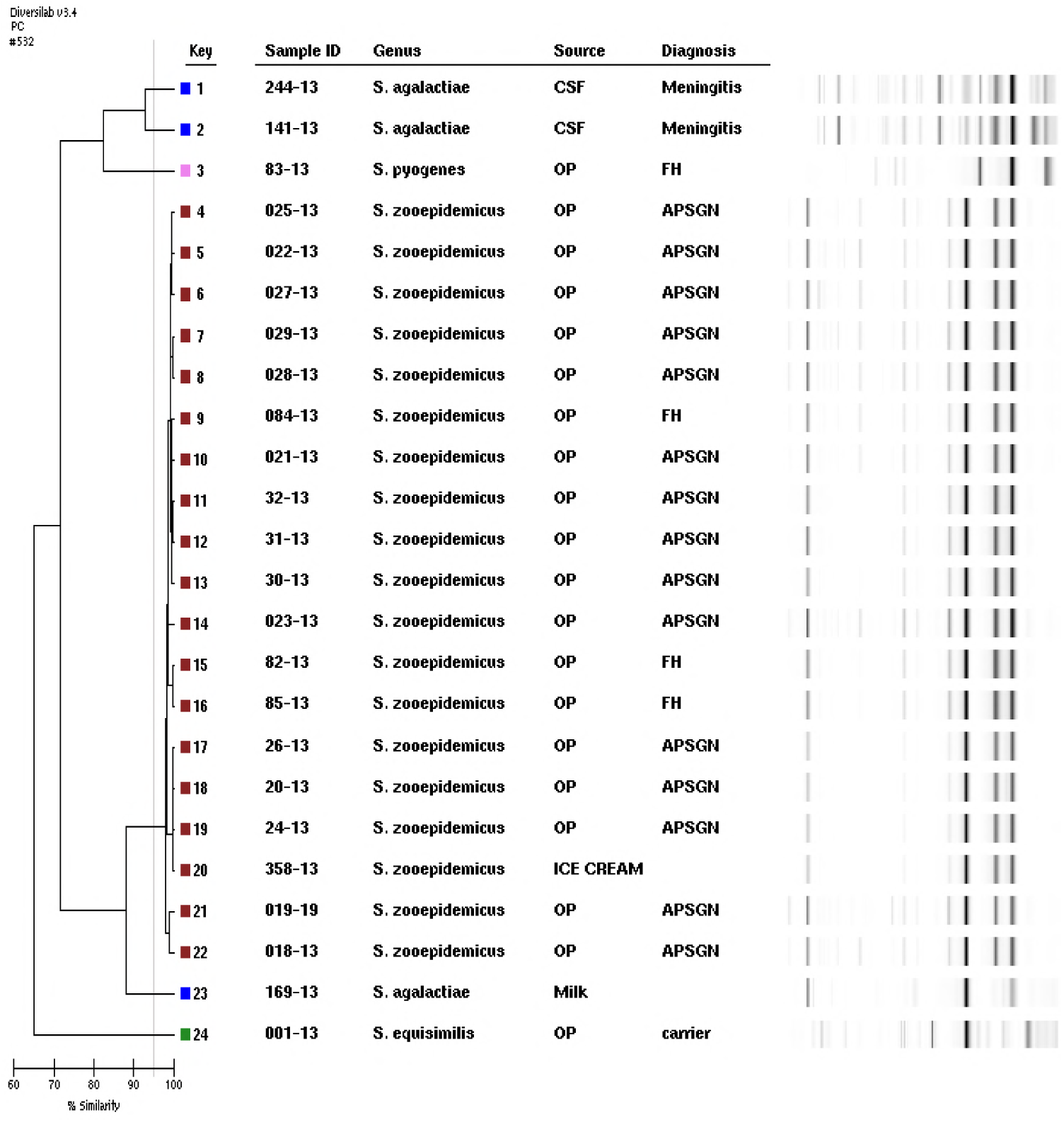
Rep-PCR dendrogram and simulated electrophoresis, strains are numbered 1 to 24, and fingerprint types (P) are color-coded. Internal control: *S. agalactiae* meningitis isolate corresponding to keys 1 and 2; *S. disgalactiae* subsp. *equisimilis* carrier isolate, key 24. *S. zooepidemicus* (outbreak strains) corresponding to keys 4-22; *S. agalactiae* isolated from milk, key 23 and *S. pyogenes emm-*type 3 oropharynx of food handler, key 3. Legend: OP, oropharynx; FH, food handler.

### Discussion

**Outbreaks** induced by emergent pathogens represent a health risk factor for a large number of individuals if quality controls in the production of milk and in the chain of milk-derived sub-products are not preserved. *S. zooepidemicus* is an emerging and opportunistic zoonotic pathogen, associated with severe and life-threatening diseases and potentially capable of triggering large glomerulonephritis outbreaks. It causes diseases in humans through the contact with infected animals (23-26) or the consumption of milk and dairy products (3,4).

**This** study discussed genetically isolated *S. zooepidemicus* associated with a large outbreak of pharyngitis and APSGN in Monte Santo de Minas, MG, Brazil. The microorganism was identified in the oropharynx of APSGN patients, food handlers and ice cream samples. All isolates involved in the outbreak amplified the same sequence of the hypervariable region of the SzP protein and were classified as SzPHV5. The isolates were categorized as clonal, due to the indistinguishable banding patterns with high similarity (> 99%) by rep-PCR, characterizing the genetic relationship between them and confirming the outbreak.

**Previous** studies report a milk-borne outbreak with 85 cases of sore throat, and later APSGN, in the town of P. Neamtz, Romania (1968), where *S. zooepidemicus* szpHV3 gene was identified in lymphnode biopsies and pharyngeal exudate of patients, carriers and in cow’s milk samples (22-27). Fifteen years later another outbreak by *S. zooepidemicus szp* HV5 took place in North Yorkshire, England (1983. Three cases of nephritis after mild upper respiratory-tract infection were identified in five members of a family running a small dairy farm; of the 12 people hospitalized, eight died (3). These outbreaks implicated non-group-A streptococci in the aetiology of APSGN.

**In** Brazil, *S. zooepidemicus szp* HV5 was responsible for a large outbreak of APSGN (1997-1998), in the city of Nova Serrana, Minas Gerais / Brazil. At that time, 133 cases of APSGN were confirmed, three patients died, seven progressed to acute renal failure, and required dialysis, and 96 patients were hospitalized. During the outbreak, 4 isolates were recovered among the patients. No isolate was recovered in the analyzed milk or cheese samples (4). The sequence described in the current study shared sequence identity with SS1215 APSGN/England/1992 (3) and *szp*5058-APSGN/Nova Serrana-MG, Brazil/1998 (22).

**The** szp gene shares structural and functional similarity with the EGA M protein, mainly with the prototype of the subfamily E *emm*-types (2, 4, 49, 60, 61 and 63). Moreover, it does not have the amino terminal regions (A and B) repeated from the N-terminal region and it does not have the C region repeated from the carboxy-terminal region (found in all EGA *emm*-types) (14,28). Considering that both species are capable of triggering glomerulonephritis and that several studies associate EGA protein M with the development of the APSGN, we can infer that rheumatoid epitopes are located in the carboxy terminal D region.

**Genotyping** is a valuable tool for epidemiological surveillance of microorganisms, leading to an efficient investigation of microbial diversity, micro-evolutionary shifts, and transmission tracking. The diversity of the SzP protein HV1-5 may establish the clonal character of the infections of the respiratory tract of horses (*29*) and may also genetically characterize strains associated with outbreaks in humans (3,22,27).

**It** was not yet possible observe any relationship between a type of SzP and a specific clinical manifestation in horses (14). However, *S. zooepidemicus* SzPHV5 is likely to present superior nephritogenic properties when compared with the other SZP protein types.

**The** SzP (M-like) is capable of inducing a specific protective immune response against infection by *S. zooepidemicus*. Specific antibody targeting SzP proteins were associated with generalized diffuse glomerulonephritis in horses (6), and were also found in the serum of patients recovering from APSGN. Serum from patients in their convalescent phase revealed the presence of antibodies highly reactive against the antigens of the isolated strains, associating them with the cause of the PSGN outbreak (22). The APSGN is widely believed to be caused when antibody–antigen immune complexes lodged in the kidney glomerulus, trigger proinflammatory immunologic processes, and produce organ injury. Many extracellular streptococcal products have been causally implicated, including streptokinase, streptococcal pyrogenic exotoxin B (SpeB, an extracellular cysteine protease), glyceraldehyde-3-phosphate dehydrogenase (GAPDH), and others. However, the absence of cysteine proteinase (SpeB) in the strain *szp*5058-APSGN/Nova Serrana-MG, Brazil/1998 contradicted the belief that cysteine proteinase could induce PSGN (30).

**This** was the third major outbreak of APSGN caused by the *S. zooepidemicus* described in the world. The isolation of *S. zooepidemicus* in an ice cream sample confirmed the control case study carried out at the time of the outbreak (2012-2013); and associated ice cream with the disease transmission chain. It is possible to infer that the epitopes of the innermost portion of the protein M EGA, SzPHV5 and SzPHV3 (D region) may be involved in the development of APSGN, and related studies should be conducted. Stricter controls and supervision in the production of milk and in the dairy by-products chain should be continuously encouraged so that consumers’ health is preserved.

